# Linking serotonergic median raphe input to dorsal CA1 with mnemonic functions

**DOI:** 10.1101/2023.09.04.556213

**Authors:** Jill Gerdey, Olivia Andrea Masseck

## Abstract

The role of serotonergic signaling within the hippocampus and its role in mnemonic function is still not well understood. We used optogenetics to specifically alter median raphe serotonin input to the dorsal CA1 subfield to investigate its role in hippocampus-dependent behavior. Unexpectedly, neither activation nor inhibition of serotonin release at dCA1 fiber terminals significantly modulated object recognition, spatial memory, or anxiety behavior. Nevertehless, we observed opposite effects of increased and decreased serotonergic input on spatial learning, and a marked reduction in the use of a direct search strategy in spatial navigation following activation of serotonergic terminals in dCA1, i.e. release of serotonin.Furthermore, since the 5-HT_1A_ receptor is one of the most abundant serotonergic receptors in CA1, we also optogenetically activated 5-HT_1A_ pathways specifically in pyrdamidal neurons of dCA1. Activation of 5-HT_1A_ signaling significantly improved spatial memory without affecting object recognition or avoidance behavior. In conclusion, our data reveal modulatory effects of serotonin specifically on the acquisition of spatial memory.

## Introduction

To date, the function of catecholaminergic input to the hippocampus has been described in detail (Kempadoo et al., 2016; Moreno-Castilla et al., 2017). However, although serotonin (5-HT) is known to be an important modulator of behavior, e.g. for behavioral adaptation, strategy optimization, cognitive flexibility, reversal learning and behavioral inhibition (Cohen et al., 2015; Cools et al., 2008; Deakin and Graeff, 1991; Lottem et al., 2018; Matias et al., 2017; Ranade and Mainen, 2009) and the serotonergic system provides a dense input to the hippocampus (Ihara et al., 1988), the detailed role of serotonin in hippocampal information processing remains elusive. Several 5-HT receptor subtypes are expressed in the hippocampal formation, but their function in mnemonic function is poorly understood. The most abundant receptor within the hippocampal formation is the 5-HT_1A_ receptor, which is expressed on pyramidal neurons and granular cells, as well as on glia (Andrade and Nicoll, 1987; Chalmers and Watson, 1991; Gross et al., 2002; Riad et al., 2000). Within the stratum lacunosum moleculare also the 5-HT_1B_ receptor is highly expressed at axon terminals from the median raphe nuclei (Winterer et al., 2011). In addition, 5-HT_2A/2C_, 5-HT_4_ and 5-HT_7_ receptors are also found within the hippocampal formation {Tanaka:2012ct}.Serotonin can potentiate excitatory synaptic input from the entorhinal cortex and promote action potential output from CA1 (Cai et al., 2013). Several studies pinpoint to an involvement of the 5-HT_4_ receptor in potentiation of the Schaffer Collateral CA3-CA1 pathway and memory function {Hagena:2017gg, Mlinar:2006bu, Teixeira:2018gw}.

Many conflicting studies have been published on the influence of serotonin on memory processing and cognition, in part because of the nonspecific, often systemic application of serotonin and ligands (a systematic review of the role of serotonin in declarative memory can be found elsewhere, for example in (Coray and Quednow, 2022). For example, Teixera et al. 2018 showed an improvement in spatial memory by optogenetically evoked terminal release of 5-HT in dorsal CA1 (Teixeira et al., 2018), while in contrast, depletion of serotonin in the hippocampus facilitated spatial learning in other studies (Gutiérrez-Guzmán et al., 2011; 2017). For instance memory consolidation in a contextual fear protocol was impaired by optogenetic stimulation of median raphe neurons (Wang et al. 2015).

In general, theta oscillations play a key role in hippocampal learning. Theta oscillations occur especially during REM sleep and exploratory behavior (e.g.: (Buzsáki et al., 1983; Jouvet, 1969; Staudigl et al., 2012)). An increase in high theta frequencies has been observed during the use of cognitive maps in the Morris-Water maze (Olvera-Cortés et al., 2004) and during memory consolidation when sharp wave ripples dominate (Buzsáki, 2015). In particular, theta is thought to promote the ability to process incoming signals (Buzsáki, 2002) and to regulate attentional processes and information processing (Nuñez and Buño, 2021). Several studies support the hypothesis that coupling of theta within the hippocampus facilitates spatial encoding (Colgin and E. I. Moser, 2009; Takahashi et al., 2014). Serotonin is known to regulate theta rhythm in the hippocampus (Kinney et al., 1995; Marrosu et al., 1996; Vertes, 2006; Vertes et al., 1994), and neuronal firing within the raphe nuclei is phase-locked to theta rhythms (Bland et al., 2016; Buzsáki, 2002; Vertes and Kocsis, 1997). The activity of GABAergic interneurons in the MR, which act directly on serotonergic output neurons, also correlates well with theta (Jelitai et al., 2021). In addition, fast and slow gamma oscillations in the hippocampus are also involved in spatial memory (Bieri et al., 2014). Systemic application of 8-OH-DPAT can affect both theta and gamma (Xu et al., 2016). We propose that serotonergic neuromodulation, particularly via pyramidal 5-HT_1A_ receptors, can modulate sensory information processing in the hippocampus and that this is critical for the interpretation and representation of sensory experiences. In order to test our hypothesis, we started to analyze which part of the serotonergic system gives rise to projections to the hippocampus in order to subsequently optogenetically stimulate or inhibit the release of serotonin in the dorsal CA1 (dCA1) and to additionally modulate 5-HT_1A_ receptor signalling in pyramidal neurons only.

## Results

### Input to the dorsal CA1 region arises mainly from the median raphe nucleus

To study the input from the raphe nuclei to the hippocampus (HPC) we used retrograde tracing in combination with immunostaining to identify serotonergic cell bodies that project to the dorsal CA1. The majority of inputs to the hippocampus arises from the median raphe nucleus (MR) (**Figure 1 A-C**). To identify the main targets of serotonergic axonal projections in dCA1 we injected a floxed tdtomato in the MR of SERT-Cre mice (**Supplementary Figure 1**) and quantified expression of terminals in the CA1 region broken down for all layers (SR, SO, SLM) (**Fig. 1D-F**). Most of the serotonergic fibers terminate in the stratum radiatum (SR) and stratum lacunosum-moleculare (SLM) of CA1 (**Figure 1F**), consistent with previous reports that also show of a laminar distribution of serotonergic fibers (Ihara et al., 1988; Varga et al., 2009; Vertes et al., 1999).

**Figure 1:**
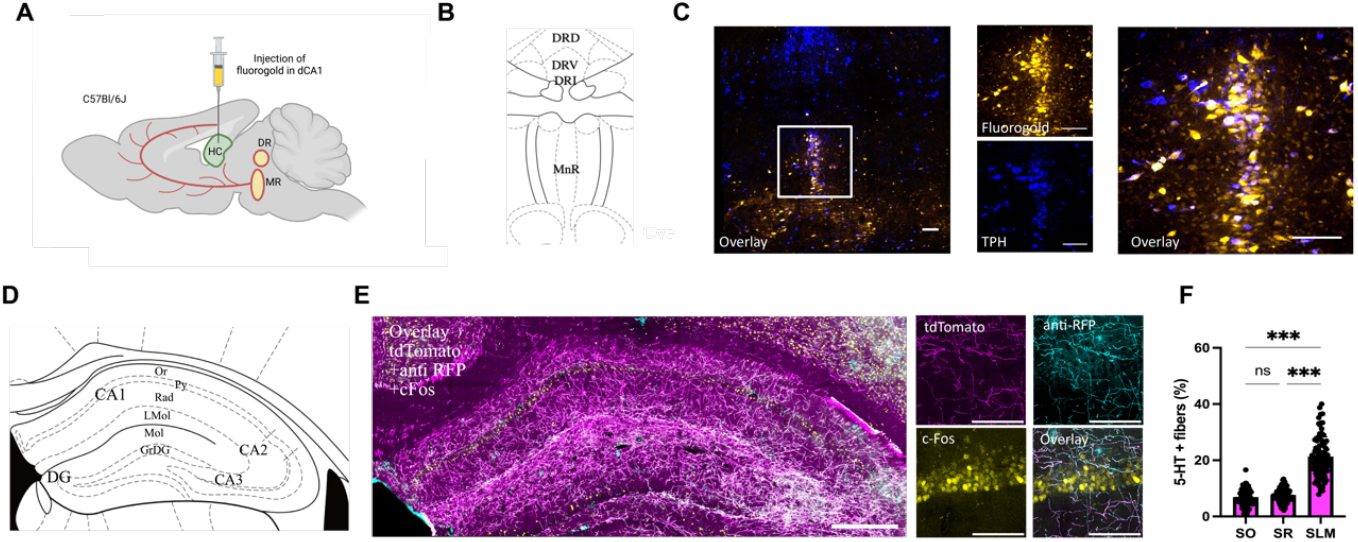
Main input to the hippocampus arises from the median raphe nucleus (MR). (**A**) Schematic drawing fluorogold injection in dCA1 of C57Bl/6J mice. (**B**) Left: Schematic overview of the raphe nuclei (RN) adapted from the mouse brain atlas of Paxinos and Franklin. (**C**) Confocal image showing coronal overview of the RN. Fluorogold labeled neurons are shown in yellow and TPH as marker for serotonergic neurons in blue. Scale bar, 100 μm. Close ups show an overview of colocalization of Fluorogold (yellow) and TPH (blue) in the MR. Scale bars, 100 μm. DRD: dorsal part of DR, DRV: ventral part of DR, DRI: interfascicular part of DR, MnR: median raphe nucleus. (**D**) Schematic overview of dorsal hippocampus with cell layers adapted from mouse brain atlas of Paxinos and Franklin. (**E**) SERT-Cre mice were injected with a floxed-tdtomato. Representative confocal image of serotonergic fiber distribution in the dorsal hippocampus. Magenta fibers express tdTomato, cyan fibers are stained against RFP and yellow cell bodies indicate c-Fos positive cells. Scale bar, 2000 μm. Close ups show an overview of colocalization of tdTomato (magenta), anti-RFP (cyan) and c-Fos (yellow) in dCA1. Scale bars, 100 μm. (**F**) 5-HT fiber density in hippocampal subregions (One -Way ANOVA,, p < 0.001, n=). SLM, stratum lacunosum-moleculare; SO, stratum oriens; SR, stratum reticulatum. White boxes indicate location of inset. DRD: dorsal part of DR, DRV: ventral part of DR, DRI: interfascicular part of DR, MnR: median raphe nucleus, TPH: tryptophan hydroxylase, RFP: red fluorescent protein, Or: oriens layer, Py: pyramidal cell layer, Rad: stratum radiatum, LMol: lacunosum moleculare layer of the hippocampus, Mol: molecular layer of the dentate gyrus, GrDG: granular layer of the dentate gyrus.

We hypothesised that 5-HT release might increase memory performance, given findings from several studies (“Constitutive and Acquired Serotonin Deficiency Alters Memory and Hippocampal Synaptic Plasticity.,” 2017; Jardin et al., 2014; Teixeira et al., 2018), whereas inhibition of serotonin release via stimulation of 5-HT_1B_ receptors at serotonergic axon terminals would impair memory performances. To test our hypothesis, we expressed either ChR2 or light-activatable 5-HT_1B_ receptors (Masseck et al., 2014; Hasegawa et al. 2017)in serotonergic neurons of the MR in SERT-Cre mice (**Supplementary Figure S2**) and stimulated MR fiber terminals in dCA1 (**Figure 2 A-F**). Additional c-Fos staining verified the functionality of the optogenetic tools applied (**Figure 2G**). As expected, an increase in c-Fos expression was observed between naïve (homecage) and no light stimulation (within the behavioural room and attached to the optical cable without light stimulation). Light activation of ChR2 at fiber terminals significantly decreased c-Fos expression in dCA1, implying a major inhibitory effect in the hippocampus. Stimulation of 5-HT_1B_ receptors at serotonergic terminals showed no significant changes in the expression of c-Fos when analysed with a 2-Way ANOVA. However, there is a clear trend towards an overall increase in cell activity. An additional t-test between no light stimulation and light stimulation shows a significant effect (two-tailed unpaired t-test: p = 0.0044). Both observations are consistent with the existing literature showing a major inhibitory effect of serotonin in the hippocampus (Corradetti et al., 1992; Varga et al., 2009) and support the functional expression of the optogenetic tools used.

**Figure 2:**
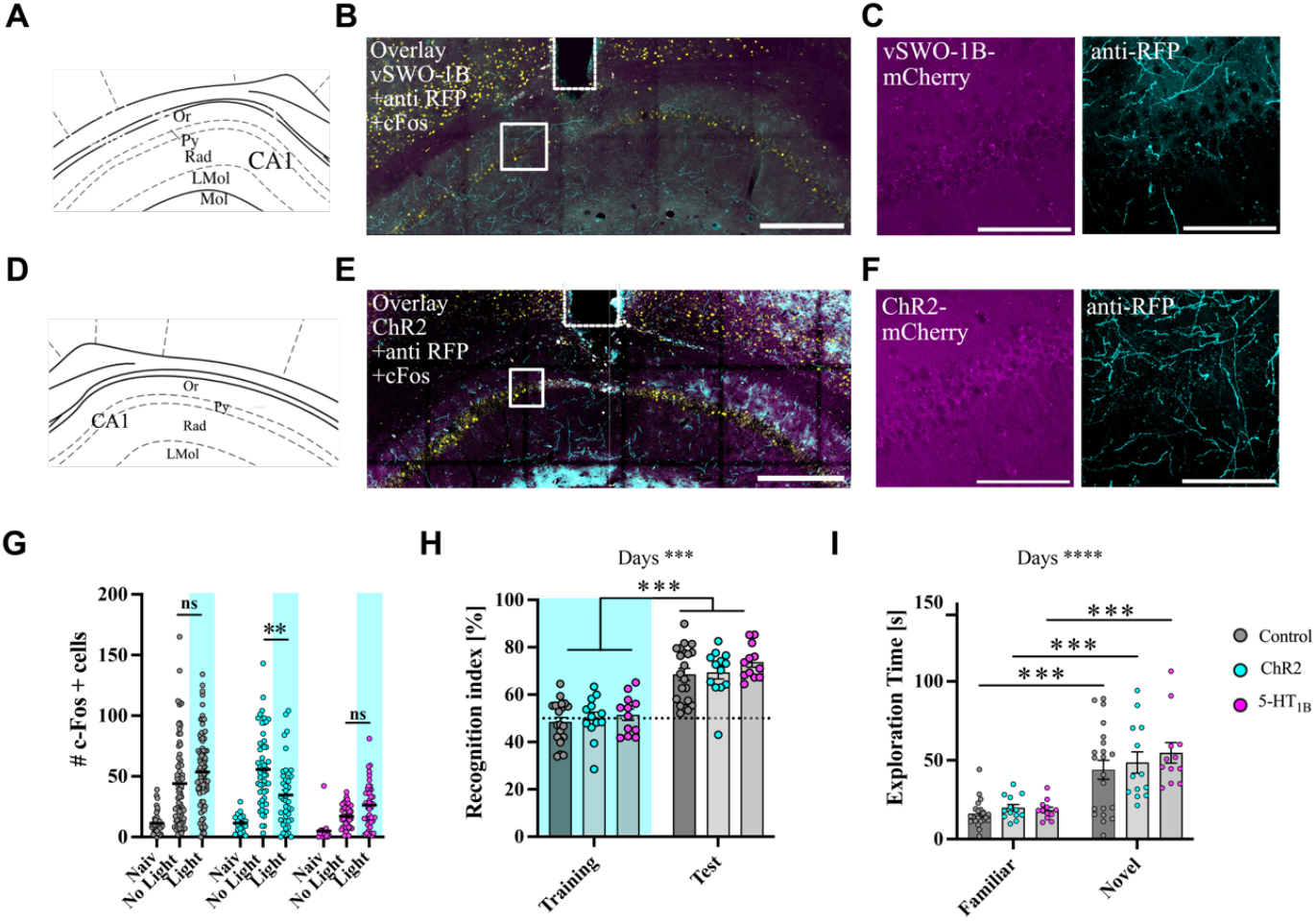
Optogenetic activation of serotonergic fibers to dCA1 do not alter object recognition. **(A)** Schematic overview of CA1 with cell layers adapted from mouse brain atlas of Paxinos and Franklin. (**B)** Representative confocal overlay image of serotonergic vSWO-5-HT_1B_ expressing fibers in CA1. Magenta fibers express vSWO-5-HT_1B_, cyan fibers are stained against RFP and yellow cell bodies indicate c-Fos positive cells. Scale bar, 2000 μm. **(C)** Close ups show an overview of colocalization of vSWO-5-HT_1B_ (magenta), anti-RFP (cyan) and c-Fos (yellow) in CA1. Scale bars, 100 μm (**D)** Schematic overview of dorsal hippocampus with cell layers adapted from mouse brain atlas of Paxinos and Franklin. (**E)** Representative confocal overlay image of distribution of serotonergic ChR2-mCherry expressing fibers in the dorsal hippocampus. Magenta fibers express ChR2, cyan fibers are stained against RFP and yellow cell bodies indicate c-Fos positive cells. Scale bar, 2000 μm. (**F)** Close ups show an overview of colocalization of ChR2 (magenta), anti-RFP (cyan) and c-Fos (yellow) in the dCA1. Scale bars, 100 μm. (**G)** Total number of c-Fos positive cells in dCA1. ChR2 activation led to a significant (p=0.003) decreased number of c-Fos positive cells by light stimulation (n=34) compared to no light (n=56) (two-way ANOVA, interaction F _(4, 405)_ = 7.801, p < 0.001, Tukey post-hoc, **p < 0.01, ***p < 0.001). n: Control: naive = 42 slices from 4 animals, no light = 71 slices from 8 animals, light = 84 slices from 9 animals, ChR2: naive = 17 slices from 2 animals, no light = 54 slices from 4 animals, light = 41 slices from 5 animals, 5-HT_1B_: naive = 18 slices from 2 animals, no light = 45 slices from 5 animals, light = 42 slices from 5 animals. (**H)** All groups showed object recognition. Neither activation of ChR2 nor 5-HT_1B_ receptor chimera showed a significant impact on object recognition (two-way mixed ANOVA, Days: F _(1, 43)_ = 111.0, p < 0.001, Tukey post hoc, ***p < 0.001, Group: F _(2, 43)_ = 1.553, p = 0.223). **(I)** In all groups, the exploration of the novel object is two times higher than for the familiar object. No difference in exploration time between the two groups could be observed (two-way mixed ANOVA, Days: F _(1, 43)_ = 76.14, p < 0.001, Bonferroni post hoc, ***p < 0.001, Group: F _(2, 43)_ = 0.8207, p = 0.447). Values represent mean ± SEM. ***p < 0.001. White boxes indicate location of inset. DRD: dorsal part of DR, DRV: ventral part of DR, DRI: interfascicular part of DR, MnR: median raphe nucleus, TPH: tryptophan hydroxylase, RFP: red fluorescent protein, Or: oriens layer, Py: pyramidal cell layer, Rad: stratum radiatum. LMol: lacunosum moleculare layer of the hippocampus.

## Behavioral significance of MR serotonergic pathways

### Object recognition task

First, we examined the behaviour of mice in a non-spatial object recognition task (NORT) after habituation to the arena animals were allowed to freely explore the open arena with two different objects. On the next day one of the objects was exchanged with a new one. Bilateral intrahippocampal light stimulation (20 Hz, 10ms) was performed during the training sessions only by which either the release or inhibition of serotonin was provoked by light stimulation. **(Fig.2 H-I)**. We did not observe any significant difference during stimulation of serotonergic fibers, neither optogenetic stimulation of serotonin release nor inhibition via light activation of chimeric 5-HT_1B_ receptors led to an impairment or improvement in object recognition (**Fig.2 H-I**). Animals in all three groups showed an increased recognition index during the test, indicating that they recognized the new object as novel. However, there was no difference for the discrimination index between the groups. An increase in exploration times supported this observation as well. The mice in each group interacted with the new object twice as long as with the familiar one. However, there is no difference in exploration time between groups **(Fig. 2I)**. Interestingly the number of visits at the new object were increase for both experimental groups during test day (**Suppl. Fig. S3A)**. However, the mean duration at the novel object was not changed (**Suppl. Fig. S3B**). Changes in the number of visits at the new object were accompanied by an increase in distance and velocity of ChR2 and 5-HT_1B_ animals (**Suppl. Fig. S3 C-D)**. During light stimulation 5-HT_1B_ mice showed an increased exploration behavior marked by a significant increase in distance travelled and velocity (**Suppl. Fig. S3 C-D)**

Although it is mainly the ventral hippocampus that is known to be involved in anxiety behavior (Jimenez et al., 2018), a recent study showed that also serotonergic input to the dorsal hippocampus promotes anxiogenic behavior (Abela et al., 2020). Therefore, we additionally investigated the effect of optogenetic stimulation in the elevated plus maze (EPM). An interaction between time and group was found for the time spent in the open arms **(Suppl. Fig. S4 A)**. The control group and 5-HT_1B_ spent significantly more time in the open arms over time, whereas no significant changes were found for the ChR2 group. The Turkey post-hoc test showed no significant effects between groups. The control group and the 5-HT_1B_ group spent more time in the open arms over time, resulting in a shorter path length. **(Suppl. Fig. S4 B-D)**. In contrast, the number of open arm entries was not affected by light stimulation **(Suppl. Fig. S4 C)**. Taken together, the data show an increase in exploration behavior of the open arms in the 5-HT_1B_ group.

The 5-HT_1A_ receptor is abundant on pyramidal neurons of CA1 and as knockout mice show an impairment in spatial as well as in memory in general (Sarnyai et al 2000), we next assessed the influence of 5-HT_1A_ signaling in memory performance. For this purpose, we expressed a light-activatable 5-HT_1A_ receptor chimera(Masseck et al., 2014; Oh et al., 2010) in pyramidal neurons of dCA1 (**Fig. 3A-C**).c-Fos stainings showed that by light stimulation the overall activity in dCA1 was decreased (**Fig. 3D**). In the light of the inhibitory action of the G_i_ coupled 5-HT_1A_ receptor this supports reports from the literature that have shown that 5-HT_1A_ receptors is the main serotonergic receptor being expressed in CA1. Also, direct stimulation of 5-HT_1A_ signaling pathways in dCA1 pyramidal neurons did not foster any differences in memory performances (**Fig. E-F**). Neither frequency, mean duration at the novel object are changed due to activation of 5-HT_1A_ signalling pathways in dCA1 pyramidal neurons, also distance and velocity are not altered **(Suppl. Fig. S5**). As 5-HT_1A_ receptor agonist can alter the anxiety behavior and 5-HT_1A_ receptor knockout mice show an increase in anxiety behavior, we also assessed anxiety behavior. Surprisingly no significant differences in anxiety or exploratory behavior could be detected due to stimulation of 5-HT_1A_ signaling pathways in pyramidal neurons of dCA1 (**Suppl. Fig. S6**). But, both control and experimental groups only rarely visited the open arms **Suppl. Fig. S6A)**. A closer look reveals that there is a tendency for light stimulation of 5-HT1A receptors in dCA1 to increase time in open arms without changing exploratory behavior, indicating a less pronounced involvement of dCA1 in anxiety behavior.

**Figure 3:**
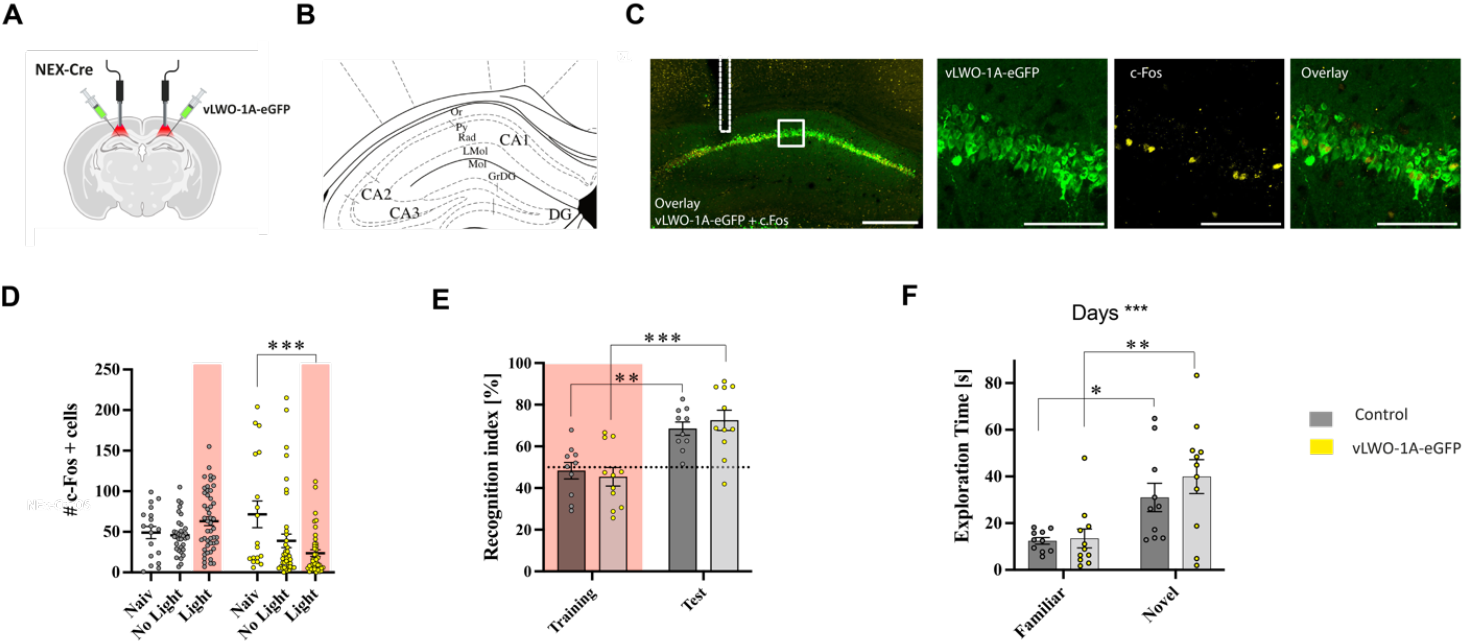
Optogenetic activation of light-activatable 5-HT_1A_ receptors in pyramidal neurons of dCA1 does not alter object recognition. **(A)** Schematic of experimental setup: viral vector coding for vLWO-5-HT_1A_-eGFP is injected into dCA1 of NEX-Cre mice. Bilateral implantation above dCA1 allowed light stimulation during behavioral tests. **(B)** Schematic overview of dHPC with cell layers adapted from the mouse brain atlas of Paxinos and Franklin. **(C)** Representative confocal image showing the expression of vLWO-5-HT_1A_-eGFP in dCA1. Green cell bodies express vLWO-5-HT_1A_-eGFP and yellow cells indicate c-Fos positive cells. White dotted line shows the placement of the optical fiber. Scale bar, 2000 μm. Close ups show an overview of colocalization of vLWO-5-HT_1A_-eGFP (green) and c-Fos (yellow) in the CA1. Scale bars, 100 μm. **(D)** Total number of counted c-Fos positive cells in dCA1 (two-way ANOVA, Interaction: F _(2, 196)_ = 8.016, p < 0.001, Turkey post-hoc, ***p < 0.001). n: Control: naive = 18 slices from 2 animals, no light = 34 slices from 4 animals, light = 48 slices from 4 animals, 5-HT_1A_: naive = 18 slices from 2 animals, no light = 42 slices from 4 animals, light = 42 slices from 5 animals.**(E)** Both groups showed object recognition, yet activation of 5-HT_1A_ receptor chimera showed no influence on object recognition when compared to control group (two-way mixed ANOVA, Days: F _(1, 19)_ = 30.99, p < 0.001, Bonferroni post-hoc, **p < 0.01, ***p < 0.001), (Groups: F (1, 19) = 0,02, p = 0.901). **(F)** In both groups the exploration of the novel object is two times higher than for the familiar object. No difference in exploration time between the two groups could be observed (two-way mixed ANOVA, Days: F _(1, 19)_ = 23.30, p < 0.001, Bonferroni post-hoc, *p < 0.05, **p < 0.01), (Groups: F (1, 19) = 0,7302, p = 0.403).Values represent mean ± SEM. *p < 0.05, **p < 0.01, ***p < 0.001.

### Spatial memory Barnes maze

To explore the role of serotonin in spatial learning we performed the Barnes maze (BM). During this behavioral assessment of hippocampus dependent spatial memory mice explore a circular platform illuminated with an aversive bright light, with 20 equidistant holes and one escape box. During 4 training sessions with 4 trials a day animals learn to find the escape boy. On day 5 and day 12 animals were challenged in a probe trial to find the target hole (escape box was removed) (**Fig. 4 A-B)**. Light stimulation was applied during all training sessions. During training, no differences were observed for optogenetic stimulation of serotonergic fibers (ChR2) or inhibition (5-HT_1B_)of serotonin release at serotonergic fiber terminals to dCA1 (**Fig. 4C-F**). During training, all groups improved their performance as indicated by a decrease in primary and escape latency **(Fig. 4C-D)**. Neither the primary nor the escape latency was significantly different between groups. This was also the case for the escape latency of the first trial **(Fig. 4E)** or the primary latency **(Suppl. Fig.7A**) of the first trial. Furthermore, the difference between escape and primary latency (**Fig. 4F**) did not differ between groups neither during all trials or the first trial **(Suppl. Fig.7B**). Also, no differences between the distance moved between the groups was evident **(Suppl. Fig.7C**). The only significant effect could be seen in the latency to reach the TH position for the ChR2 activated group on the test days (**Fig. 4G**). They needed less time to reach the TH position on day 12 than on day 5. However, the amount of time spent in the TH quadrant was the same for all groups on both days of testing (**Fig. 4H**). Neither the primary latency nor the escape latency differed significantly between groups, nor did the frequency and number of errors differ for all groups tested. This was also the case for the escape latency (**Fig. 4E**) or the primary latency (**Suppl. Fig. 7A**) of the first trial. Furthermore, the difference between escape and primary latency (**Fig. 4F**) did not differ between groups during all trials or during the first trial (**Suppl. Fig. 7B**). There were also no differences in the distance traveled between groups (**Suppl. Fig. 7C**). The time spent around TH was reduced in all groups on day 12 compared to day 5 (**Fig. 4I**). No differences in the frequency of TH visits, number of errors or preference index were observed (**Suppl. Fig. 7D-F**). However, the 5-HT_1B_ group has a tendency to be superior to the ChR2 groups in almost all parameters under observation. Next, the strategy for finding the escape box was evaluated. Modulation of serotonergic input to dCA1 did not appear to have a direct effect on escape behavior during the classical Barnes maze paradigm, but the choice of strategies differed (**Fig. 4J-H**). Animals in which serotonin release was stimulated during training (ChR2) used the direct strategy significantly less than the control group on day 3 and on test day 5 **(Fig. 4J-H**). Accordingly, animals in which serotonin release was evoked during training used a mixed strategy more often than the other groups (**Supplementary Figure 7E**).

**Figure 4:**
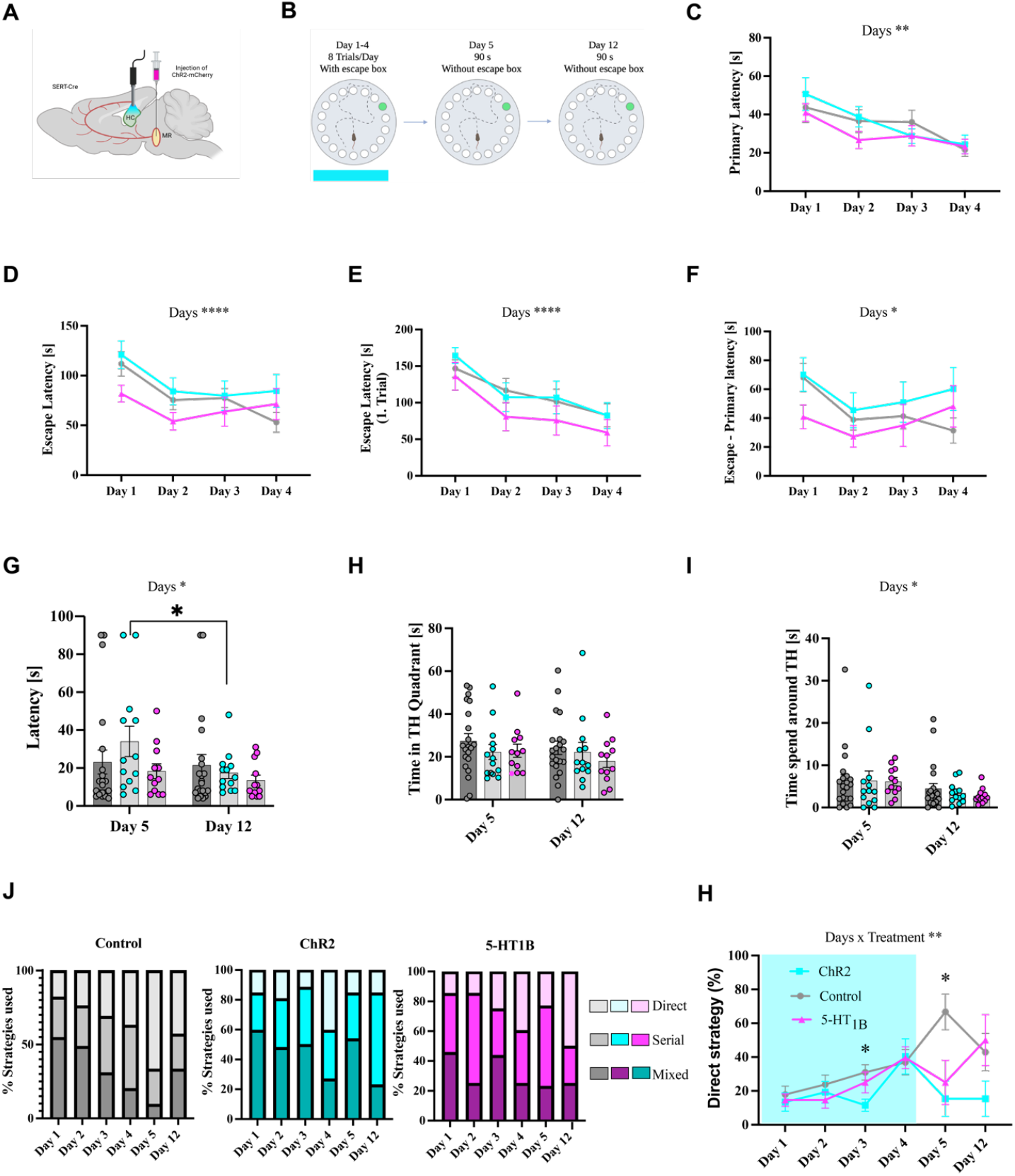
Optogenetic activation of serotonergic fibers to dCA1 changes strategies utilized in the Barnes Maze. **(A)** Schematic of experimental setup: viral vector for either vSWO-5-HT_1B_ or ChR2d is injected into dCA1 of. Bilateral implantation above dCA1 allowed light stimulation during Barnes Maze. **(B)** Experimental protocol: Four consecutive days of learning (with light stimulation) followed by two test trials without escape box with one week in between (without light stimulation). **(C)** All three groups showed improved primary latency over days however no difference between activation of ChR2 (cyan), activation of vSWO-5-HT_1B_ receptor chimera (magenta) and control group (grey), (two-way mixed ANOVA, Days: F (2.031, 87.33) = 9.541, p < 0.001, Bonferroni post-hoc, Control: Day 3 vs Day 4 *p < 0.05, ChR2: Day 2 vs Day 4 *p < 0.05, 5-HT1B: Day 1 vs Day 4 *p < 0.05). **(D)** Groups did not differ in escape latency and only control animals improved their performance from day 1 to day 4 (two-way mixed ANOVA, Days: F (2.363, 101.6) = 10.17, p < 0.001, Bonferroni post-hoc, control: Day 1 vs Day 2 **p < 0.01, Day 1 vs Day 4 ***p < 0.001, 5-HT1B: Day 1 vs Day 2 *p < 0.05). **(E)** Escape latency for the first trial of training day was not different between groups. A significant difference for main effect of days was observed. (two-way mixed ANOVA, Days: F (2.682, 115.3) = 12.79, p < 0.001, Bonferroni post-hoc, control: Day 1 vs Day 4 *p < 0.05, ChR2: Day 1 vs Day 4 *p < 0.05). **(G)** Latency to reach TH position was decreased for activation by ChR2 from day 5 to day 12 (two-way mixed ANOVA, Days: F (1, 43) = 5.306, p = 0.026, Bonferroni post-hoc, *p < 0.05). **(F)** Only the control group showed significant decrease in the difference between escape and primary latency over days (two-way mixed ANOVA, Days: F (2.348, 101.0) = 3.947, p = 0.017, Bonferroni post-hoc, Control: Day 1 vs Day 2 **p < 0.01, Day 1 vs Day 4 *p < 0.05). **(H)** No difference in spending time in TH quadrant (two-way mixed ANOVA, Days x Group: F (2, 43) = 0.5534, p = 0.579). **(I)** A significant difference for the main effect of days was seen for the time spent around TH (two-way mixed ANOVA, Days: F (1, 43) = 5.337, p = 0.026, Bonferroni post-hoc showed no significant comparisons). **(J)** Search strategies for experimental and control groups. **(H):** ChR2 group used the direct strategy significant less on day 3 and day 5. (two-way mixed ANOVA, Days x Group: F (10, 215) = 2.635, p = 0.005, Bonferroni post hoc, Control: Day 1 vs Day 5 **p < 0.01, Day 2 vs Day 5 **p < 0.01, Day 3 vs Day 5 *p < 0.05). Values represent mean ± SEM. *p < 0.05, **p < 0.01, ***p < 0.001.

Next, we explored the influence of 5-HT_1A_ signaling on spatial navigation (**Fig. 5 A-B**). Mice in which we stimulated 5-HT_1A_ receptor chimeras in dCA1 pyramidal neurons showed improved performance during training. Both groups learned the location of the escape box as evidenced by decreased primary and escape latencies over time (**Fig. 5C-F**). The 5-HT_1A_ receptor chimera activated group showed improved performance during training compared to the control group (**Fig. 5D-F**). Notably, not all mice entered the escape box on their first visit, so the time it took them to enter the escape box on their first visit (primary latency) was also plotted (**Fig. 5C**). Primary latency decreased during the training days for both groups, although it did not change between the two groups. In addition, the primary escape latency of the first trial during the day was significantly different between groups **(Supplementary figure S8A)**. 5-HT_1A_-stimulated mice were faster to escape from the open platform (Figure 5D). Since the escape latency is an average of four trials, it is not a reflection of how much the mice had remembered from the previous day. Indeed, the improvement was still observed when only the first trial of the day was plotted (**Fig. 5E**). To analyze whether there was a difference in the time spent on the open platform, the difference between escape latency and primary latency was calculated (**Fig. 5F**). The 5-HT_1A_ group spent less time on the open platform than the control group, and both groups spent less time on the open platform over days. The 5-HT_1A_ mice took less time to enter the box even when only the first runs of the day were considered (**Supplementary Figure S8B**). Thus, the acquisition of spatial learning is enhanced by acute stimulation of 5-HT_1A_ signaling in dCA1 pyramidal neurons.

**Figure 5:**
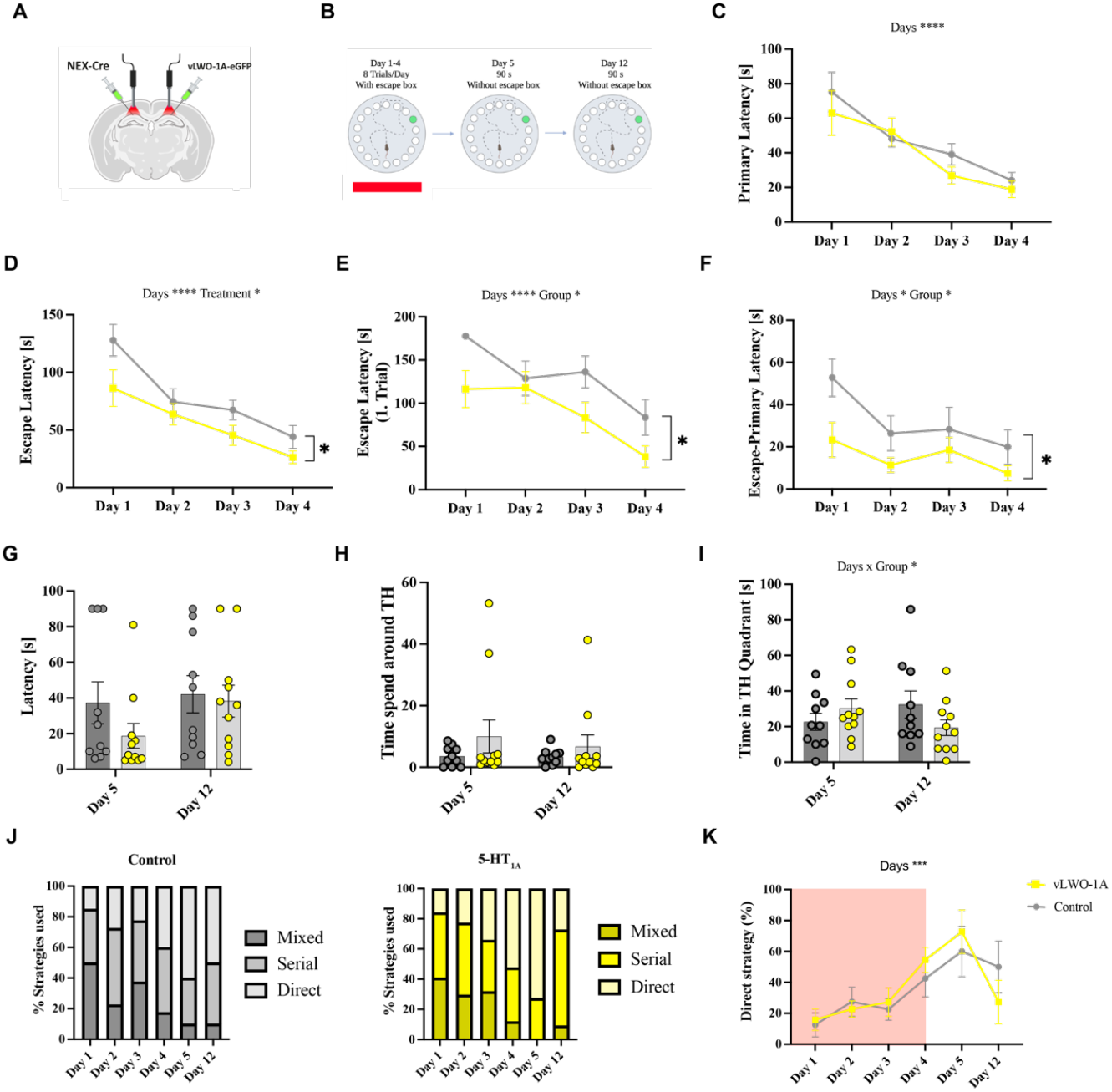
Acute stimulation of 5-HT1A signaling pathways in dCA1 enhances spatial learning. **(A)** Experimental setup: vLWO-5-HT_1A_-eGFP is expressed in pyramidal neurons of dCA1. Bilateral implantation above dCA1 allowed light stimulation during behavioral tests. **(B)** Experimental protocol: Four consecutive days of learning (with light stimulation) followed by two test trials without escape box with one week in between (without light stimulation). **(C)** Latency to the first visit of escape box was not different between control (grey) and 5-HT_1A_ group (yellow), nevertheless both groups improved their performance during training (two-way mixed ANOVA, Days: F _(1.804, 34.28)_ = 18.62, p < 0.001, Bonferroni, Control: Day 1 vs Day 4 **p < 0.01, Day 2 vs Day 4 *p < 0.05, 5-HT_1A_: Day 1 vs Day 2 *p < 0.05, Day 1 vs Day 4 *p < 0.05, Day 2 vs Day 4 **p < 0.01, Group: F _(1, 19)_ = 0.7412, p = 0.400). **(D)** Both groups showed an improved performance during training, whereas 5-HT_1A_ activation caused a faster escape latency (two-way mixed ANOVA, Days: F _(2.382, 45.25)_ = 23.97, p < 0.001, Group: F _(1, 19)_ = 4.541, p = 0.046, Bonferroni post-hoc did not show significant multiple comparisons). **(E)** 5-HT_1A_ activated animals needed less time to escape from the open platform on the first trial of the day. All animals improved performance during training (two-way mixed ANOVA, Days: F _(2.678, 50.88)_ = 10.45, p < 0.001, Bonferroni post-hoc, Control: Day 1 vs Day 4 **p < 0.01, 5-HT_1A_: Day 1 vs Day 4 **p < 0.01, Day 2 vs Day 4 **p < 0.01), Group: F _(1, 19)_ = 7.730, p = 0.012, Bonferroni post-hoc did not detect differences between groups). **(F)** Significant group effect for difference between escape and primary latency. 5-HT_1A_ activation leads to a faster entering of escape box. Both groups improved performance over days (two-way mixed ANOVA, Days: F _(2.049, 38.93)_ = 4.804, p = 0.013, Bonferroni post-hoc, Control: Day 1 vs Day 2 *p < 0.05, Day 1 vs Day 3 *p < 0.05, Day 1 vs Day 4 ***p < 0.001, 5-HT_1A_: Day 1 vs Day 4 **p < 0.01, Day 2 vs Day 4 **p < 0.01, Group: F _(1, 19)_ = 7.082, p = 0.015, Bonferroni post-hoc did not detect differences between groups). **(G)** Latency to reach TH position on test day was not significantly different (two-way mixed ANOVA, Days x Group: F _(1, 19)_ = 0.6734, p = 0.422). **(H)** No significant difference for time spent around TH during test days (two-way mixed ANOVA, Days x Group: F _(1, 19)_ = 0.5988, p = 0.449). **(I)** A Days x Group interaction effect was observed for time spent in quadrant, however Bonferroni post-hoc analysis didn’t show differences (two-way mixed ANOVA, Days x Group: F _(1, 19)_ = 5.871, p = 0.026, Bonferroni post-hoc did not detect differences between groups or days). **(J)**: Graphical overview of search strategies for control group. **(K)**: Graphical overview of search strategies for 5-HT_1A_ group. **(L)**: Both groups learned the position of the escape box over days and used more the direct strategy (two-way mixed ANOVA, Days: F _(3,441, 65,37)_ = 7,361, p < 0.001, Bonferroni post-hoc, Control: Day 1 vs Day 4 *p < 0.05, 5-HT_1A_: Day 1 vs Day 4 *p < 0.05, Day 1 vs Day 5 *p < 0.05, Day 2 vs Day 5 *p < 0.05, Group: F _(1, 19)_ = 0.009788, p = 0.922, Bonferroni post-hoc did not detect significant differences). Values represent mean ± SEM. *p < 0.05, **p < 0.01, ***p < 0.001.

By removing the escape box on the fifth day, it is possible to analyze how well the mice remembered the location of the TH. This test day was repeated one week later to examine long-term memory. No significant differences were found between the groups in the latency to reach the TH (**Figure 5G**) or in the time spent around the TH (**Fig. 5H**). A significant Days x Group interaction was calculated for the time spent in the TH quadrant (**Fig. 5I**), while Bonferroni post hoc did not show significant differences. This may indicate that the effect of acute light stimulation is transient and important only during memory acquisition.

Spatial strategies were recorded in order to investigate different types of search behavior. In general spatial learning is characterized by a decrease in the mixed strategy (**Supplementary Figure S8G**) and an increase in the direct strategy over time (**Figure 5J-K**). This was observed for both groups (**Fig. 5J-K**) and no difference in the use of the direct strategy was found between the groups (**Fig. 5K**).

In conclusion, these results demonstrate an improved acute performance during spatial learning by light activation of the 5-HT_1A_ receptor chimera in dCA1 pyramidal neurons, which did not alter the animals’ search strategy.

## Discussion

In agreement with many other studies (Freund, 1992; Ihara et al., 1988; Jacobs et al., 1974) our tracing experiments confirmed that the main serotonergic input originates from the MR. Only parts of the caudal dorsal raphe nucleus innervate the dorsal hippocampus (Vertes, 1991; Vertes et al., 1999). A clear distinction between target areas of the MR and the DR, which mainly innervate the ventral hippocampus is present. Serotonergic innervation showed a laminar distribution with the highest input to the stratum lacunosum moleculare (Varga et al., 2009; Vertes et al., 1999).

Neither serotonin release, inhibition of serotonin release, nor activation of 5-HT_1A_ pathways in dCA1 affected object recognition. There is currently an ongoing debate as to whether object recognition is mediated by the dorsal hippocampus at all, as several studies have shown that lesions within the hippocampus only affect spatial memory and not object recognition (Duva et al., 1997). The NORT used in our study did not have a spatial component, which may explain why we did not observe any effects on object memory. Local infusion of a 5-HT_1B_ agonist in the dorsal hippocampus induced a neophobic response and a decrease in locomotor activity leading to avoidance in an object recognition task (Buhot and Naïli, 1995). However, these effects occurred only when the animals were first exposed to the novel environment. In all of our behavioral experiments, the animals were habituated to the arena, which may explain why we did not find a decrease in exploratory behavior. Besides, our studies selectively stimulated 5-HT_1B_ receptors on serotonergic terminals (i.e. decreaseing serotonin release), whereas local infusion of a 5-HT_1B_ agonist also targets cholinergic axon terminals of septohippocampal where it inhibits the release of acetylcholine release in addition. In the same study intrahippocampal injections of 8-OHDPAT (5-HT_1A_ and 5-HT_7_ agonist), decreased object exploration and habituation in rats ((Buhot and Naïli, 1995).We did not see any significant changes induced by 5-HT_1A_ receptor stimulation on object recognition or exploratory behavior, suggesting an exclusive role of 5-HT_1A_ in spatial memory.

No change in anxiety behavior was observed in any of the groups. One reason for this may be that the animals were already handled and habituated to the procedures when the EPM was performed, raising the possibility that innate anxiety behavior is already attenuated in all experimental animals, making it difficult to assess anxiety behavior. Furthermore, anxiety behavior is classically assigned to the ventral hippocampus (Jimenez et al., 2018). A recent study could show an involvement of MR serotonergic input to the dorsal hippocampus on approach-avoidance behavior in female mice (Abela et al., 2020), but failed to show behavioral changes in the EPM. This is consistent with our data in male mice.

Our data suggest that 5-HT_1A_ receptor signaling on pyramidal neurons in dCA1 play a main role in the acquisition of spatial memory. This is well in line with spatial learning deficits observed in 5-HT_1A_ knockout mice (Sarnyai et al., 2000) and with intrahippocampal infusions of 8-OHDPAT that provoked an increased responsiveness to a spatial changes of a familiar object (Buhot and Naïli, 1995). Improved spatial learning might pinpoint to a prominent engagement of slow neuromodulatory action of 5-HT_1A_ receptor activation at pyramidal neurons in spatial memory acquisition,whereas probably fast direct synaptic transmission via glutamate/serotonin cotransmission at hippocampal interneurons serve other functions (Varga et al., 2009).

We could not replicate the findings of Teixera et al 2018, where optogenetic stimulation of serotonergic fiber terminals in dCA1, i.e. serotonin release increased spatial memory and inhibition of serotonin release impaired spatial memory (Teixeira et al., 2018). This may be due to methodological differences: In our experiments, we used SERT-Cre mice and expressed ChR2 via a viral-based approach only in MR serotonergic neurons, whereas Teixera used a transgenic ePet1 mouse line that expressed ChR2 in all serotonergic neurons, including the dorsal raphe nucleus (DR). As a consequence terminal stimulation in dCA1 will also activate serotonin release from the DR. In addition, the number of targeted neurons in ePet (Scott et al., 2005) and SERT (Zhuang et al., 2005) mice differ from each other (Cardozo Pinto et al., 2019). The important role of region- and input-specific modulation of neuralmodulatory signals is demonstrated by these contrasting results.

Theta oscillations within the hippocampal network are thought to be important regulators in the emergence and coding properties of place and grid cells (Buzsáki, 2002; E. I. Moser and M.-B. Moser, 2008; O’Keefe and Recce, 1993), which are the building blocks for spatial learning. Activity of place cells is presumed to be an allocentric representation of space, necessary for spatial memory (Dupret et al., 2010; O’Keefe, 1976; Wilson and McNaughton, 1993).

Several studies implicating a modulatory role of 5-HT on hippocampal oscillations, I,e, theta, gamma and ripples. MR neuron activity is tightly coupled to theta oscillations, and an increase in theta amplitude and synchronization, as required for spatial learning, only occurs when serotonergic input is withdrawn. In contrast, activation of MR neurons abolishes hippocampal ripples (Wang et al. 2015; Xu et al., 2016). Our behavioral data partially support the hypothesis that 5-HT suppression may facilitate the emergence of theta and ripples, as indicated by a trend toward improved acquisition in the 5-HT_1B_ group compared to a deterioration in ChR2 mice.

We suggest that activation of somatic 5-HT_1A_ receptors on pyramidal neurons, in concert with inhibition of interneurons (Royer et al. 2012), may be critical for controlling firing patterns in pyramidal neurons. In our experiments, inhibition of a large population of dCA1 pyramidal neurons by light activation of 5-HT_1A_ receptors could support population coding by reducing redundancy in the network (Abbott and Dayan, 1999; Zohary et al., 1994), potentially leading to improved spatial learning. Hence, mnemonic function may critically depend on the level and pattern of 5-HT1A receptor expression within the hippocampal circuitry.

## Acknowledgements

O.A.M received funding for this project by the DFG (SFB 874- Projektnummer 122679504) pAAV9-EF1a-double-floxed-ChR2(H134R)-mCherry-WPRE-HGHpA (Addgene number #20297-AAV9) was a gift from Karl Deisseroth and pAAV9-FLEX-tdTomato (Addgene number #28306-AAV9) a gift from Edward Boyden. We thank Prof. K. Nave and Dr. S. Goebbels for kindly providing NEX-Cre mice and Xiaoxi Zhaung for SERT-Cre mice.

## Author contribution

J. G. and O.A.M designed and analysed experiments. J.B. performed behavioral experiments and immunohistochemical stainings. J.B and O.A.M wrote the paper.

## Methods

### Animals

All experiments were performed in accordance with the guidelines and approval of the Senator für Gesundheit, Frauen und Verbraucherschutz der Freien Hansestadt Bremen (TV 147). Adult (8-16 weeks old) male Nex-Cre, SERT-Cre and wild-type C57BL/6J mice were used in the following experiments. All animals were housed in standard individually ventilated cages (IVC, Zoonlab) with controlled temperature (22°C □ 2°C) and humidity (50% □ 5%). All cages were equipped with houses, tunnels, and bedding. Mice were given food and water ad libitum and maintained on a 12/12-hour reverse light cycle (lights on at 22:00). All behavioral experiments were performed in the dark. Mice were housed in groups until surgery. After surgery, animals with implants were housed individually to prevent damage to the implants. All animals were bred in-house.

### NEX-Cre

The NEX-Cre transgenic mouse line (Goebbels et al. 2006) was used to express a vLWO 5 HT1A mCherry chimera in dCA1 pyramidal neurons. The NEX-Cre line expresses Cre recombinase under the control of the NEX gene (Goebbels et al., 2006). Only heterozygous animals were used in this study.

### SERT-Cre

The SERT-Cre line was used to express ChR2 and vSWO-5-HT1B in serotonergic neurons (Zhuang et al., 2005). Mice were bred in-house using breeding pairs purchased from The Jackson Laboratory (B6.129(Cg)-Slc6a4tm1(cre)Xz/J, stock #014554).

### Virus injection and optical fiber implantation

#### Optical implants

Custom-made optical implants (Berg et al. 2020) were used for light stimulation during behavioral testing. A glass optical fiber (200 μm in diameter; Thorlabs, FP200URT), cut to a length of 2.5 cm, was inserted into a ceramic ferrule (Thorlabs, CFLC230-10) and secured with superglue. After fixation, the glass fiber was cut near the convex end of the ceramic fiber. To improve the light output, the end of the fiber was polished with decreasing grits of fiber polishing paper (30 μm, 6 μm, 3 μm, and 1mm). Finally, the fiber was cut to the desired length with a diamond knife (Thorlabs, S90R). The hippocampal implants had a final length of 1.5 mm. The light output of the implants was measured with a photometer (Thorlabs, PM100D) for verification. The light output of the implants used in this work ranged from 2 to 4.5 mW/mm^2^. For in vivo light simulation, the mouse implants were connected to a Y-shaped optical cable (Prizmatix) attached to an LED light box (Prizmatix).

### Stereotactic surgery

All mice were at least 8 weeks old before virus injection. Mice were initially anesthetized with 5% isoflurane and then maintained under 1.5%-2% isoflurane (cp-pharma, Isoflurane CP®). All animals received 5 mg/kg carprofen (Zoetis, Rimadyl®) for analgesia. Once the mouse was in deep anesthesia (no toe reflex), it was fixed in a stereotactic frame (Stoelting Co.) with an integrated heating pad. The skull was oriented in all directions. The mice were continuously treated with ophthalmic ointment (Bepanthen) to prevent dehydration of the eyes during surgery. A scalp incision was then made from anterior to posterior. The skull was roughened with 37.5% phosphoric acid (Kerr, OptiBond Set) for a maximum of 15 seconds to facilitate subsequent fixation of the implant. Burr holes for virus or tracer injection were then drilled into the skull above the regions of interest (Coordinates for virus injection: MRN: AP: -4.6, ML: 0, DV: -4.5; dCA1: AP: -2.1, ML: ±1.5, DV: -1.5 coordinated to bregma).

A volume of 1 μl virus was injected at each injection site. When only performing injections, the incision was sewn shut (SMI, 191050) after the viral injection to conclude the surgery. For implantations, the skull surface was further treated with a thin layer of primer and adhesive from an OptiBond™ FL kit (Kerr) which were cured with UV light (850 W/cm^2^). Optical fibers were fixed into a custom-made holder for the stereotactic frame in order to correctly insert the implants into the drilled holes. The optical fibers were lowered to the desired position (dCA1: AP: -2.1, ML: +-1.5, DV: -1.4) and fixed with dental cement (Geiz Dental GC 2278). To prevent inflammation, the skin around the surgical area was treated with iodine ointment (Betaisodona^®^). Implanted animals were separated into cages and single-housed to prevent damage to the optical fibers. Animals were given three weeks for recovery and virus expression before starting with behavioral experiments.

### Experimental Groups

For behavioral tests, folwoing cuvadeno associated viruses were used: pAAV9-EF1a-double-floxed-ChR2(H134R)-mCherry-WPRE-HGHpA (Addgene number #20297-AAV9) pAAV9-FLEX-tdTomato (Addgene number #28306-AAV9), pAAV9-vLWO-5HT_1A_-eGFP (custom made) and pAAV9-SWO-5HT_1B_-mCherry (custom made). All mice were implanted bilateral targeting the dCA1.

For retrograde tracing experiments, 1 μl of 2% Fluorogold (dissolved in NaCl) was injected bilaterally into the dCA1 (AP: -2.1, ML: +-1.5, DV: -1.5). For immunostaining of Fluorogold injections, mice were perfused 2–5 days after injections and expression was verified with fluorescence microscopy.

### Behavioral Tests

All behavioral experiments were performed in the active phase (dark-phase) of the animals. Testing was conducted 3 weeks after surgery. Thirty minutes before testing, the animals were brought to a small hallway in front of the behavioral room. Experiments were recorded and analyzed using EthoVision XT (Noldus). Light stimulation was performed with a Prizmatix LED box (STSI blue and orange-red). Continuous red light (610 nm; ∼2-4 mW output) was used to stimulate vLWO-5-HT_1A_ and the corresponding controls (Masseck et al., 2014). The vSWO-5HT_1B_ chimera and related controls were stimulated with continuous blue light (460 nm; ∼2-4 mW output; (Masseck et al., 2014), while ChR2 and related controls were stimulated with 10 ms pulses of blue light (460 nm; ∼2-4 mW output) at a frequency of 20 Hz (Teixeira et al., 2018). After each trial, all mazes were cleaned with 70% ethanol.

Behavioral tests were performed in the following order: Barnes maze test, novel object recognition task, elevated plus maze. Tests were performed at one-week intervals. After behavioral testing, mice were transcardially perfused and c-Fos staining was performed.

### Barnes Maze

The Barnes Maze (BM) is a behavioral test that utilizes the natural escape behavior of rodents to study their spatial learning abilities. This behavioral test is less stressful than the Morris water maze (Morris et al., 1982; Sunyer et al., 2007). It consists of a circular platform (92 cm in diameter) with 20 identical holes (5 cm in diameter) surrounding the outer circumference of the platform. All holes were closed except for one where an escape box was placed underneath. Visual cues (a triangle, circle, and cross) were placed on the three walls of the behavioral chamber and remained consistent throughout training and testing. To further stimulate escape behavior, the room was illuminated with bright LED lights (1000 lux). Mice were placed in the escape box for two minutes prior to the first habituation trial. Each trial began by placing the mouse in a start box in the center of the maze to ensure that the mouse started in a random direction. Mice were trained on four consecutive days, each day with four trials at 15-minute intervals. If the mice were unable to enter the escape box within three minutes, they were gently redirected. After each trial, they were allowed to spend one minute in the escape box before being returned to their home cage. On the fifth day, the test day, the escape box was removed and the hole was closed. Mouse behavior was recorded for 90 seconds. After one week (day 12), a test day without the escape box was repeated to examine long-term memory. In all experiments, light stimulation was only applied during training trials.

Spatial strategies were categorized into three different strategies: mixed, serial, or direct. A mixed strategy was used when the mouse randomly crossed the center of the maze without a clear strategy. The serial strategy was used when a mouse systematically searched from hole to hole. If the animal went directly to the area of the escape box (within two holes of the box) and entered the escape box, a direct strategy was used (Harrison et al., 2006). One problem with the Barnes maze is that some mice do not enter the box on their first visit. This problem was observed in previous studies and solved by additionally recording the primary latency (time required for the first visit of the escape box; (Harrison et al., 2006). In addition, several other parameters were recorded (using EthoVisionXT) during training trials, such as primary latency (time to first encounter the target), escape latency (time to enter the escape box), and path length (total distance traveled). Parameters recorded in the test trial included latency to reach the target hole (TH), time spent in the quadrant of the TH, time spent near the TH, frequency of TH visits, number of errors, preference index (mean of TH visits/mean of all other hole visits), and total distance traveled.

#### Novel Object Recognition Task

The Novel Object Recognition Task (NORT) assesses the ability of rodents to recognize novel objects. It was first described by Ennaceur and Delacour (1988) and takes advantage of rodents’ natural preference for novelty. The test was conducted in a square open field arena (50×50×40 cm). The animals were first habituated to the open field arena for ten minutes on two consecutive days. On the third day, two identical objects (250 ml bottles filled with sand) were presented to the animals and they were allowed to investigate them for 10 minutes. One hour later, one object was replaced with a novel object (a tower made of Duplo blocks). The new object was different in shape, color, and structure, but had similar dimensions. Positioning of the objects was counterbalanced across animals to avoid bias. Light stimulation was provided during training, when the identical objects were presented. The minimum amount of light required for the software to track the mouse was used during experiments.

For the NORT, mice were habituated to the open field arena for ten minutes on two consecutive days. On the third day, the animals were presented with two identical objects and given ten minutes to explore. One hour later, one of the objects was replaced with a novel object and the mice were recorded for another 10 minutes. Light stimulation was applied during the training trials.

For behavioral analysis, the time spent exploring the objects with the snout (within a 2 cm area around the object) and the frequency of interactions with the objects were recorded. The objects were specifically chosen to prevent the animals from sitting on them, which would have been erroneously counted as exploratory behavior. To quantify object recognition, a recognition index was calculated for each animal. The recognition index reflects the exploration of the familiar and the novel object compared to the total exploration.

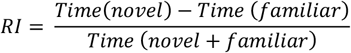

### Elevated Plus Maze

The Elevated Plus Maze (EPM) is a common behavioral test used to study anxiety behavior in rodents. It is a plus-shaped maze consisting of two open and two closed arms. The test relies on the natural aversion that mice have to open and elevated areas, as well as their natural impulsive exploratory behavior in novel environments (Komada et al., 2008). Each arm had a total length of 33.5 cm, a width of 5 cm, and closed arms had walls 17 cm high. The maze was 43 cm above the floor. The anxiety of the mice was enhanced by a bright LED light (900 lux). At the beginning of each trial, mice were placed in the center of the maze facing the same open arm. The mice were given five minutes to explore the maze. Light stimulation was interspersed with light-off periods. Time spent in the center and open arms, frequency of open arm visits, and total distance traveled were recorded.

### Immunohistochemistry and fluorescence microscopy

#### c-Fos Analysis

In this study, c-Fos was used as a marker of neuronal activity. c-Fos is an immediate early gene that is expressed in neurons after stimulation. Transcription initiation of the gene occurs immediately after stimulation and results in the gene encoding the nuclear protein Fos. c-Fos has a peak two hours after induction of gene transcription (for review, see Harris, 1998). For c-Fos analysis, mice in three different groups were treated 90 minutes prior to perfusion. One group was perfused directly from the home cage without a 90-minute delay (naive). The second group was connected to the light cable without light stimulation. And the third group was connected to the light cable with appropriate light stimulation. Mice in groups two and three were brought to the behavioral room, remained in their home cage, and were connected to the light cable for five minutes. After a 90-minute delay, they were transcardially perfused.

### Perfusion and Staining Protocol

To harvest the brain for immunohistochemical staining, mice were anesthetized with an intraperitoneal (ip) injection of 130 mg/kg ketamine and 10 mg/kg xylazine. After achieving deep anesthesia, they were transcardially perfused with 1x phosphate buffer saline (PBS) followed by 4% paraformaldehyde (PFA). After fixation, the brain was carefully removed and fixed in 4% PFA at 4°C overnight. The next day, the brain was stored in 30% sucrose until sectioning.

45 □m Coronal slices of the raphe nuclei and hippocampus were sectioned using a cryostat (Thermo Fisher Scientific) and collected in 1x PBS. To avoid non-specific antibody binding, sections were blocked with 10% normal donkey serum (NDS) and PBS containing 0.3% Triton X-100 (PBS-T) for 1.5 hours at room temperature (RT). Primary antibodies were diluted with 10% NDS in 0.3% PBS-T for 48 hours at 4°C. After another wash step (3x in PBS-T), the sections were incubated with secondary antibodies diluted with 10% NDS in 0.3% PBS-T for four hours at RT. Again, the sections were washed with 1x PBS to remove excess antibody solution. Finally, the sections were mounted on Superfrost™ slides (Thermo Scientific) and coverslipped with a mounting medium containing DAPI (ROTI® Mount FluorCare with DAPI, Carl Roth GmbH) to counterstain nuclei. Slides were stored at 4°C until fluorescence microscopy. Fluorogold-injected brains were harvested after 2-5 days without treatment and raphe slices were stained against tryptophan hydroxylase (TPH), which catalyzes the conversion of L-tryptophan to L-5-hydroxy-tryptophan and is commonly used for specific staining of serotonergic neurons (Haycock et al., 2002). The excitation wavelength of Fluorogold is in the UV spectrum with an emission of 536 nm. For better visualization of mCherry-labeled fibers within the dCA1, brain slices were stained with red fluorescent protein (RFP).

### List of Antibodies

Primary antibodies

▪ sheep-anti-TPH (MBS502127, MyBioSource, 1:1000)
▪ rabbit-anti-c-Fos (226008, Synaptic systems, 1:1000)
▪ goat-anti RFP (BYT-ORB182397, Biozol, 1:500)

Secondary antibodies for C57BL/6J: injected with Fluorogold

▪ Alexa568-donkey-anti-sheep (A-21099, Thermo Fisher 1:500)

Secondary antibodies for Nex-Cre: virally injected with vLWO-1A-eGFP and d-flox-tdTomato

▪ DyLight550-donkey-anti-rabbit (Sa5-10039, Thermo Fisher, 1:500)
▪ Alexa647-goat-anti-rabbit (711-605-152, Jackson ImmunoResearch, 1:500)

Secondary antibodies for SERT-Cre: virally injected with d-flox-ChR2-mCherry, vSWO-1B-mCherry and d-flox-tdTomato

▪ Alexa488-donkey-anti-sheep (A-11015, Thermo Fisher, 1:500)
▪ DyLight405-donkey-anti-goat (705-475-003, Jackson ImmunoResearch, 1:500)
▪ Alexa647-donkey-anti-rabbit (711-605-152, Jackson ImmunoResearch, 1:500)

### Fluorescence Microscopy

Images were captured using an LSM880 (Carl Zeiss) confocal microscope with Zen software. Sections with fluorogold were imaged using a Leica SP5 confocal microscope. Images were taken at either 20x, 25x, or 40x magnification and then processed in ImageJ. For c-Fos analysis of dCA, images were taken at 20x magnification in a two-by-five tile with 6 to 8 layers deep. Five to ten sections from each animal were analyzed and c-Fos quantified using ImageJ. The total amount of c-Fos positive cells within the dCA1 was used as an indicator of neuronal activity. Mice that did not express virus or lost the implant during testing were excluded from analysis.

### Data Analysis

All behavioral data were recorded using EthoVision XT software from Noldus. Data acquisition was performed automatically from recorded videos. Statistical analysis and graph generation were done using GraphPad Prism. Confocal images were processed for brightness and contrast using ImageJ, and quantification of c-Fos was performed using ImageJ’s pixel analyzer plugin.

### Statistical Procedures

Statistical tests were performed using GraphPad Prism9 software. For data comparing more than three groups, analysis of variance (ANOVA) was performed to test for statistically significant results. One-way ANOVA tests whether or not the means of independent sample groups are different. One-way ANOVA assumes normality and homoscedasticity of the data sets. The normality of the data was confirmed by the Shapiro-Wilk test. Homogeneity of variances between groups was tested using Bartlett’s test. For normally distributed and equal variances, a one-way analysis of variance (ANOVA) was performed. When the ANOVA showed a significant result, the Tukey post hoc test was used to show significant differences between groups. Non-parametric unpaired data were analyzed by Kruskal-Wallis ANOVA followed by Dunn’s test.

Mixed ANOVA followed by Bonferroni post hoc test was used for data comparing the behavior of consecutive days, with one between-subject variable (treatment) and one within-subject variable (days). Greenhouse-Geisser correction was used for repeated measures with more than two within-subject observations to correct for violation of the assumption.

Ordinary two-way ANOVA followed by Tukey’s post hoc test was used for c-Fos analysis. However, two-way ANOVA is considered robust to violations of normality, and non-normally distributed data were not tolerated. For all analyses, values are reported as mean ± SEM, and the significance threshold was set at ∗p < 0.05, ∗∗p < 0.01, and ∗∗∗p < 0.001.

### Declaration of generative AI and AI-assisted technologies in the writing process

During the preparation of this work the author(s) used DeepLWrite] in order to improve language and readability. After using this tool/service, the author(s) reviewed and edited the content as needed and take(s) full responsibility for the content of the publication.

