## Supplementary figures for "Linking serotonergic median raphe input to dorsal CA1 with mnemonic functions"

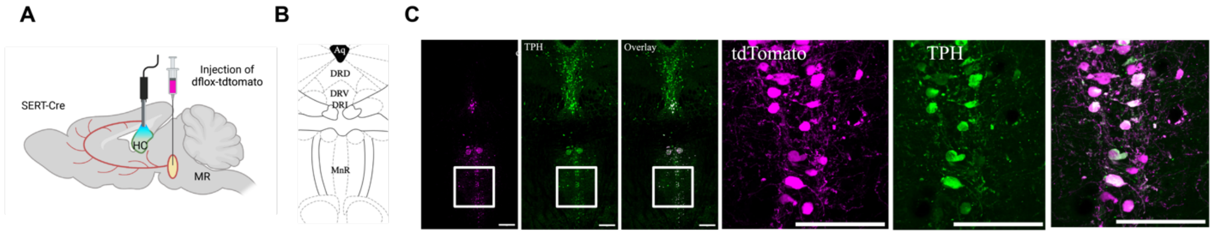


**Supplementary Figure S1: Injection of dflox-tdtomato in the MR of SERT-Cre mice.** (**A**) Schematic drawing of dflox-tdtomtao injection in the median raphe nucleus of SERT-Cre mice. (**B**) Schematic overview of raphe nuclei adapted from mouse brain atlas of Paxinos and Franklin. (**C**) Representative confocal image of a virus injection into the MR. Magenta cell bodies express tdTomato and green cells are stained with TPH. The overlay indicates the co-expression of tdTomato expressing cells and serotonergic neurons. Scale bars, 200 μm. Close ups show the localization of tdTomato (magenta) and TPH (green). Scale bars, 100 μm.)

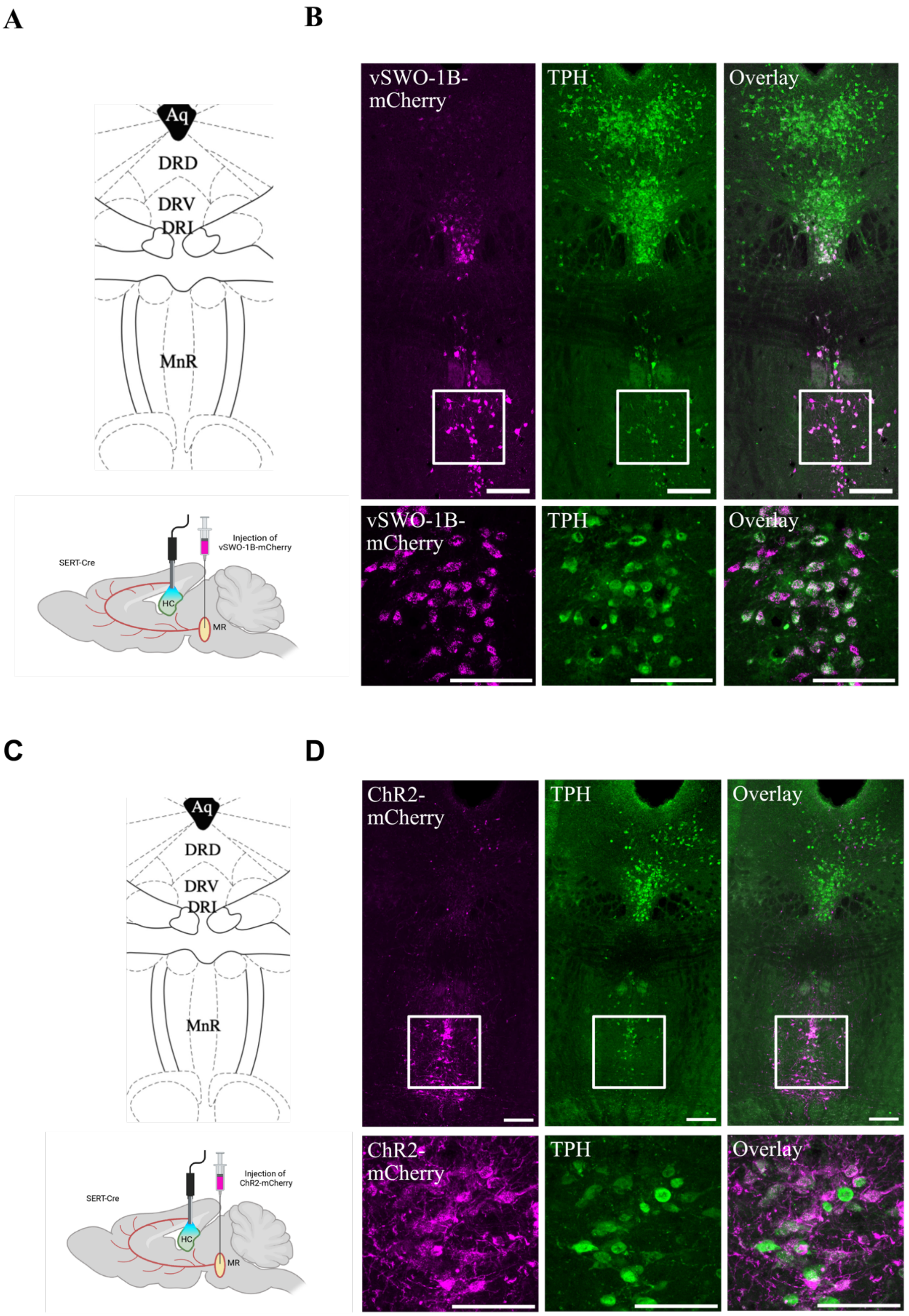


**Supplementary Figure S2: Expression of optogenetic tools in MR. (A) (C)** Schematic overview of raphe nuclei adapted from mouse brain atlas of Paxinos and Franklin Diagram of vSWO-1B virus injection in the MR. (**B)** Representative confocal image of virus injection into the MR. Magenta cell bodies express vSWO-5-HT$_{1B}$ and green cells are stained against TPH. The Overlay indicates the co-expression of vSWO-5-HT$_{1B}$ receptor chimera expressing cells and serotonergic neurons. Scale bars, 200 μm. Close ups show localization of vSWO-5-HT$_{1B}$ (magenta) and TPH (blue). Scale bars, 100 μm.**(C)** Schematic overview of raphe nuclei adapted from mouse brain atlas of Paxinos and Franklin Diagram of ChR2-mCherry virus injection in the MR. (**B)** Representative confocal image of virus injection into the MRN. Magenta cell bodies express ChR2 and green cells are stained with TPH. The Overlay indicate the co-expression of ChR2 expressing cells and serotonergic neurons. Scale bars, 200 μm. (**C)** Close ups show localization of ChR2 (magenta) and TPH (blue). Scale bars, 100 μm.

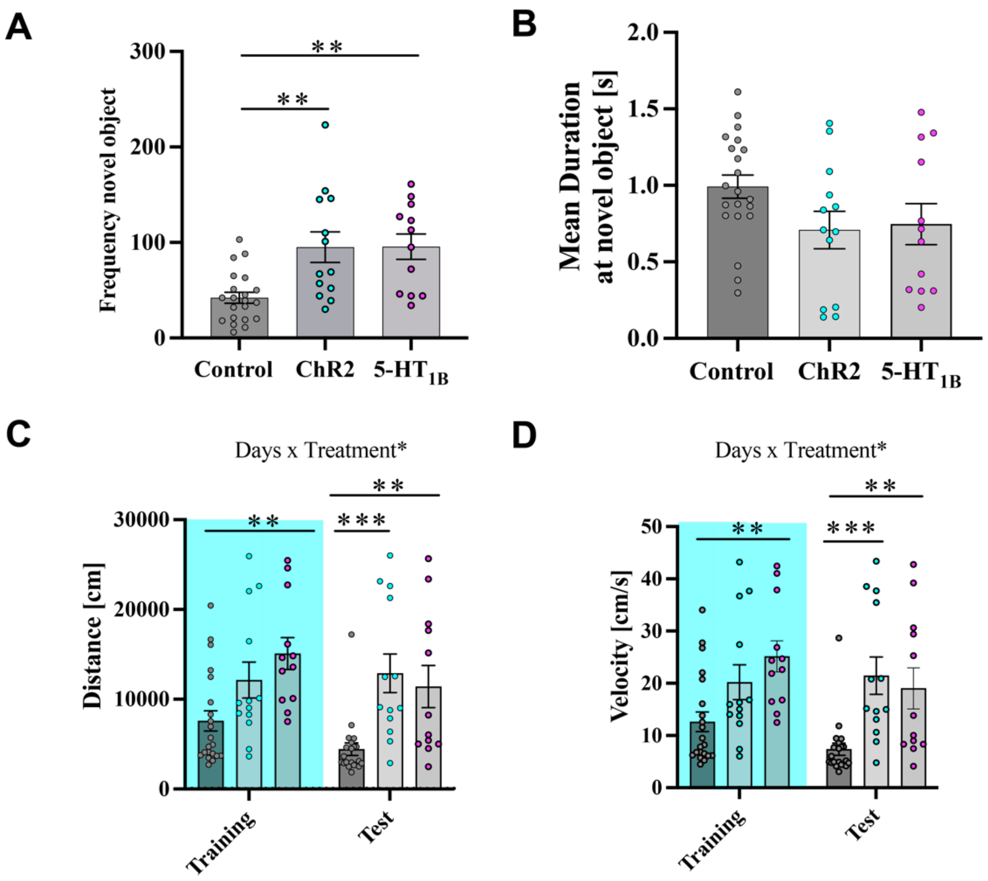


**Supplementary Figure S3: Analysis of additional parameters in the object recognition test. (A)** Frequency at the novel object. Mean control =42.05 ±5.776, ChR2= 95.15 ±16, 5-HT1B=95.58 ± 13.25.Brown_Forsythe and Welch ANOVA F(2,26.62)=7.5, **= 0.0026) (**B)** Mean duration at the novel object, no significant differences. Mean control =0.99 ±0.075, ChR2= 0.71 ±0.12, 5-HT1B=0.74 ± 0.13. One-way ANOVA, F (2,43 )=2.461, p=0.0973. **(C)** Distance traveled in cm. During the training session light stimulated 5-HT1B animals showed an increased distanced moved. During test day both experimental groups had a signigicant increased distance in comparison to the control group. Two-way ANOVA, Days x Treatment: Treatment: F(1,43)= 7.63. p=0.0084. Bonferroni multiple comparison test. Training: mean control =7588 ±1136, ChR2= 12137 ±2001, 5-HT1B=15097 ± 1776, Test: mean control =4434±703, ChR2= 12866 ±2138, 5-HT1B=11410 ± 2355. **(D)** During the training session light stimulated 5-HT1B animals showed an increased velocity in comparison to control. On test day both experimental groups had a significant increased velocity. Two-way ANOVA, Days x Treatment: Treatment: F(2,43)= 8.5. p=0.008. Bonferroni multiple comparison test. Training: mean control =12.65 ±1.89, ChR2= 20.2 ±3.33, 5-HT1B=25.17 ± 2.96, Test: mean control =7.39±1.17, ChR2= 21.48 ±3.5, 5-HT1B=19.05 ±3.93.n= 21 control mice, n=13 ChR2 , n=12 5-$HT_{1B}$. Values represent mean ± SEM. **p < 0.05. ***p < 0.001.

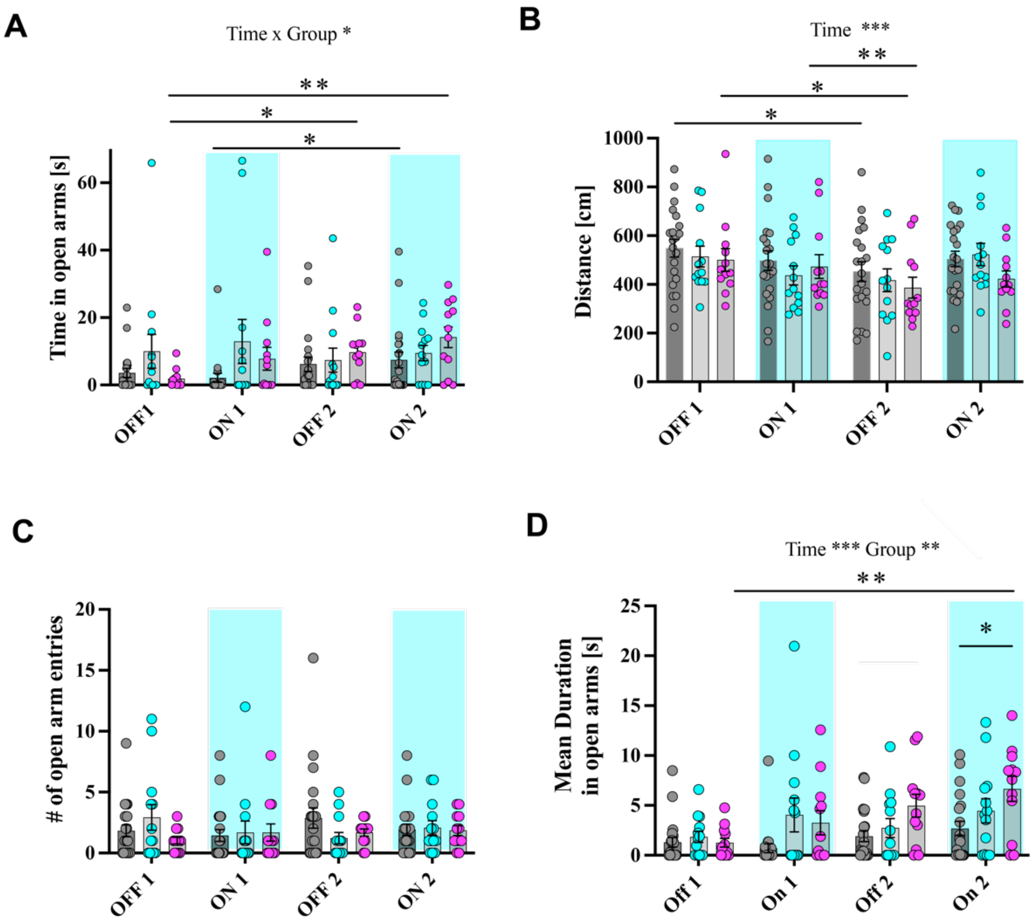


**Supplementary Figure S4: Optogenetic stimulation of ChR2 and 5-$HT_{1B}$ do not influence anxiety behavior**. **(A)** Control and 5-$HT_{1B}$ group increased time in open arms during EPM trial, whereas time in open arms for ChR2 group stays the same (two-way mixed ANOVA, Time x Group: $F_{(6, 129)} = 2.198$, $p = 0.047$, Bonferroni post-hoc *$p < 0.05$, **$p < 0.01$). **(B)** Distance decreased over time for control and 5-$HT_{1B}$ activated group (two-way mixed ANOVA, Time: $F_{(2.250, 96.75)} = 8.177$, $p < 0.001$, Bonferroni post-hoc, *$p < 0.05$, **$p < 0.01$). **(C)** Frequency of open arm entries was not altered by light stimulation (two-way mixed ANOVA, Time x Group: $F_{(6, 129)} = 1.608$, $p = 0.150$). **(D)** 5-$HT_{1B}$ group stayed longer in open arms during second light phase (two-way mixed ANOVA, Time: $F_{(2.559, 110.0)} = 7.556$, $p < 0.001$, Group: $F_{(2, 43)} = 5.567$, $p = 0.007$, Bonferroni post hoc, *$p < 0.05$, **$p < 0.01$). Values represent mean ± SEM. *$p < 0.05$, **$p < 0.01$.

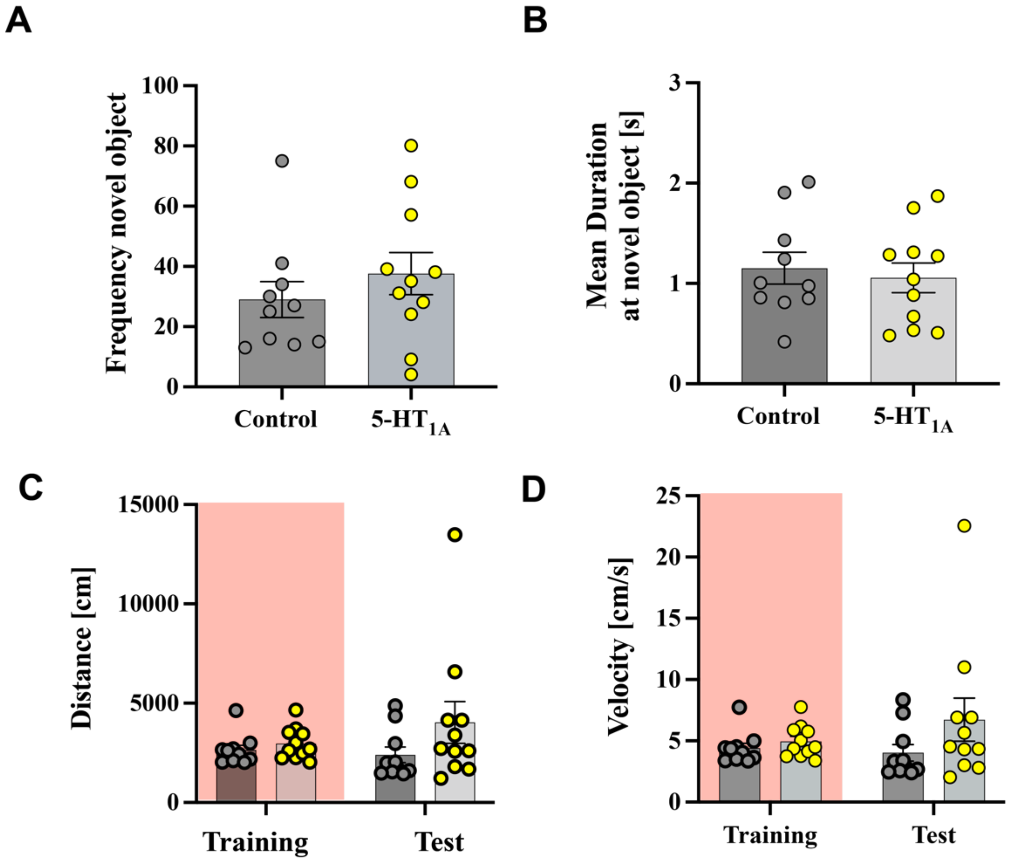


**Supplementary Figure S5: Analysis of additional parameters in the object recognition test during 5-HT$_{1A}$ stimulation. (A)** Frequency at the novel object. Mean control =29 ±5.9, 5-HT1A=37.55 ± 6.99. unpaired t-test, p= 0.3675 (**B)** Mean duration at the novel object, no significant differences. Mean control =1.15±0.159, 5-HT$_{1A}$=1.056 ± 0.14 unpaired t-test, p= 0.66 **(C)** Distance traveled in cm. During training and test session no differences between the two groups are seen. Two-way ANOVA, Days x Treatment: Treatment: F(1,19)= 2.37 p=0.1405, Days: F(1,19)=0.51, p =0.48. **(D)** During training and test session light stimulated 5-HT$_{1A}$ animals showed no differences in velocity in comparison to control (Two-way ANOVA).Test: mean control 4.33±0.4, 5-HT$_{1A}$= 4.95±0.399, Test: mean control= 4.023±0.67, 5-HT1B =6.7±1.7. n= 11 control mice, n=11 5-HT$_{1A}$. Values represent mean ± SEM.

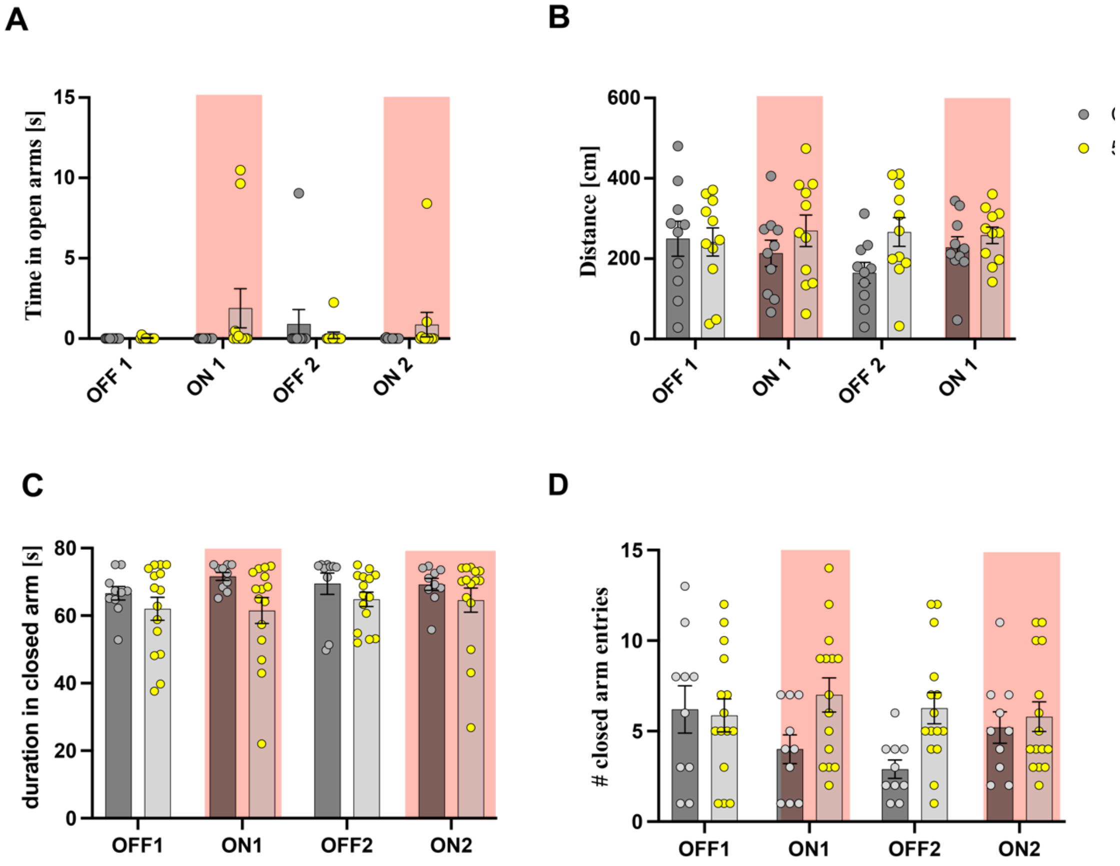


**Supplementary Figure S6: Optogenetic stimulation of 5-$HT_{1A}$ do not influence anxiety behavior. (A)** Time in open arms did not differ between groups. All mice spent only little time in open arms (two-way mixed ANOVA, Days x Group: $F_{(3, 57)} = 1.822$, $p = 0.153$) **(B)** Traveled distance was the same in both groups (two-way mixed ANOVA, Days x Group: $F_{(3, 57)} = 1.453$, $p = 0.237$). **(C)** Duration in closed arms was neither changed by light stimulation or between groups (two-way ANOVA, Inetraction: $F(3,78)=0.5843$) **(D)** Entries in closed arms is not changed. (two-way ANOVA, Interaction: $F(3,76)=1.455$) Values represent mean ± SEM.

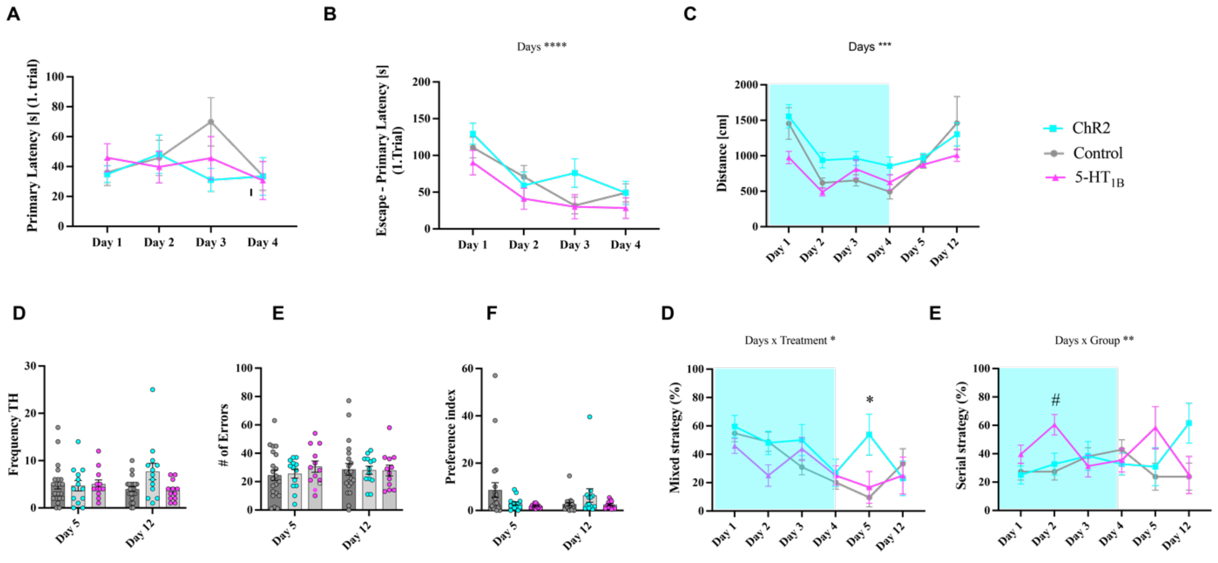


**Supplementary figure S7: Optogenetic activation of serotonergic fibers to dCA1 in the Barnes Maze. (A)** No difference was seen for primary latency of only first trial of the day (two-way mixed ANOVA, Days x Group: $F_{(6, 129)} = 1.160$, $p = 0.332$). **(B)** Groups needed less time from primary to escape latency over days (two-way mixed ANOVA, Days: $F_{(2.410, 103.6)} = 15.84$, $p < 0.001$, Bonferroni post hoc, Control: Day 1 vs Day 3 ***$p < 0.001$, Day 1 vs Day 4 *$p < 0.05$, ChR2: Day 1 vs Day 4 **$p < 0.01$, 5-$HT_{1B}$: Day 1 vs Day 3 *$p < 0.05$). **(C)** Only control and 5-$HT_{1B}$ group decreased path length over time (two-way mixed ANOVA, Days: $F_{(2.234, 96.06)} = 9.325$, $p < 0.001$, Bonferroni post hoc, Control: Day 1 vs Day 2 **$p < 0.01$, Day 1 vs Day 3 *$p < 0.05$, Day 1 vs Day 4 **$p < 0.01$, 5-$HT_{1B}$: Day 1 vs Day 2 **$p < 0.01$, Day 2 vs Day 5 **$p < 0.01$, Day 2 vs Day 12 **$p < 0.01$). **(D)** No significant differences for number of visits of TH location (two-way mixed ANOVA, Days x Group: $F_{(2, 43)} = 2.787$, $p = 0.073$). **(E)** Number of errors did not differ during groups or days (two-way mixed ANOVA, Days x Group: $F_{(2, 43)} = 1.380$, $p = 0.263$). **(F)** Preference index did not differ between groups or days (two-way mixed ANOVA, Days x Group: $F_{(2, 43)} = 3.047$, $p = 0.058$). **(G)** Usage of mixed strategy was increased for ChR2 on day 5 (two-way mixed ANOVA, Days x Group: $F_{(10, 215)} = 2.220$, $p = 0.018$, Bonferroni post hoc, *$p < 0.05$ Control: Day 1 vs Day 4 **$p < 0.01$, Day 1 vs Day 5 **$p < 0.01$, Day 2 vs Day 4 *$p < 0.05$, Day 2 vs Day 5 **$p < 0.01$). **(H)** Usage of serial strategy was increased for 5-$HT_{1B}$ on day 2 compared to control group (two-way mixed ANOVA, Days x Group: $F_{(10, 215)} = 2.948$, $p = 0.002$, Bonferroni post hoc, #$p < 0.05$ between ChR2 and 5-$HT_{1B}$, 5-$HT_{1B}$: Day 2 vs Day 3 **$p < 0.01$). Values represent mean ± SEM. *$p < 0.05$, **$p < 0.01$, ***$p < 0.001$. * indicate significant difference between control and ChR2, # significant difference between 5-$HT_{1B}$ and control group.

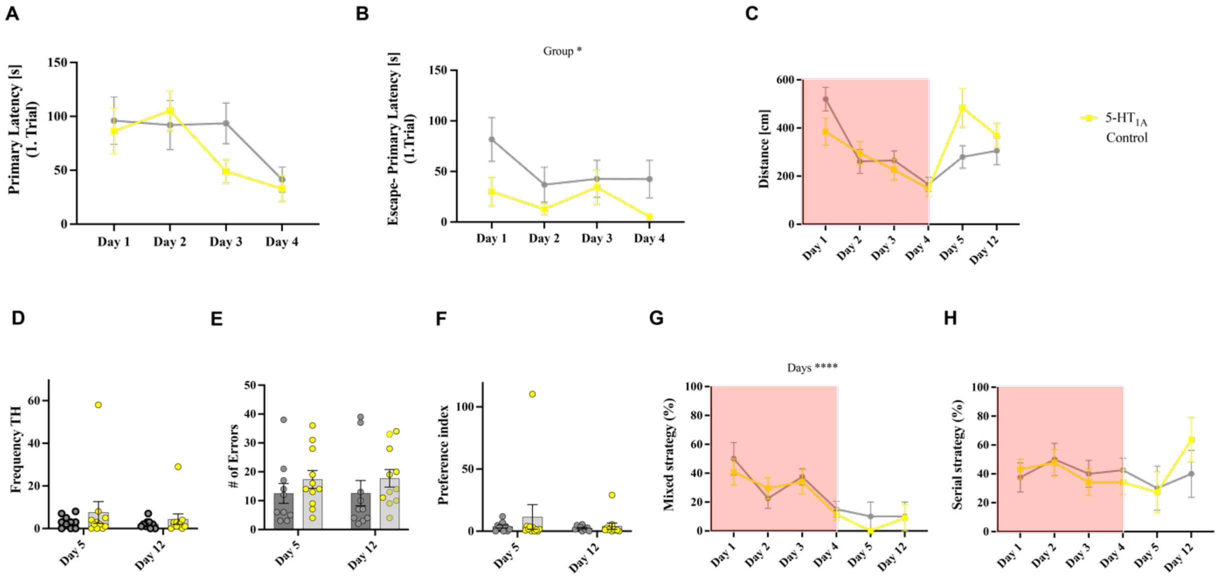


**Supplementary figure S8: Optogenetic activation of 5-HT$_{1A}$ receptors in pyramidal neurons of dCA1 in the Barnes Maze (A)** No difference was seen for primary latency of only the first trial of the day (two-way mixed ANOVA, Days: $F_{(2.559, 110.0)} = 7.556$, $p = 0.005$, Bonferroni post hoc, 5-HT$_{1A}$: Day 2 vs Day 3 *$p < 0.05$, Day 2 vs Day 4 *$p < 0.05$). **(B)** Significant group effect for escape-primary latency for the first trial of the day (two-way mixed ANOVA, Group: $F_{(1, 19)} = 7.714$, $p = 0.012$, Bonferroni post hoc did not find significant multiple comparisons). **(C)** Main effect of days for traveled distance, but no difference between the two groups (two-way mixed ANOVA, Days: $F_{(2.548, 48.42)} = 5.473$, $p = 0.004$, Bonferroni post hoc, Control: Day 1 vs Day 3 *$p < 0.05$, Day 1 vs Day 4 *$p < 0.05$). **(D)** No significant differences for number of visits of TH location (two-way mixed ANOVA, Days x Group: $F_{(1, 19)} = 0.3371$, $p = 0.568$). **(E)** Number of errors did not differ during group or days (two-way mixed ANOVA, Days x Group: $F_{(1, 19)} = 0.009$, $p = 0.926$). **(F)** Preference index did not differ between group or days (two-way mixed ANOVA, Group x Days: $F_{(1, 19)} = 1.074$, $p = 0.313$). **(G)** Usage of mixed strategy decreased over days (two-way mixed ANOVA, Days: $F_{(3.928, 74.62)} = 8.562$, $p < 0.001$, Bonferroni post hoc, 5-HT$_{1A}$: Day 1 vs Day 5 *$p < 0.05$, Day 2 vs Day 5 *$p < 0.05$, Day 3 vs Day 5 *$p < 0.05$). **(H)** Usage of serial strategy did not differ between groups (two-way mixed ANOVA, Days x Group: $F_{(5, 95)} = 0.5874$, $p = 0.710$). Values represent mean ± SEM. *$p < 0.05$, **$p < 0.01$, ***$p < 0.001$.
